# The chromosome-scale genome assembly of *Jasminum sambac* var. *unifoliatum* provide insights into the formation of floral fragrance

**DOI:** 10.1101/2022.01.25.477387

**Authors:** Chengzhe Zhou, Chen Zhu, Caiyun Tian, Siyi Xie, Kai Xu, Linjie Huang, Shengjing Wen, Cheng Zhang, Zhongxiong Lai, Yuling Lin, Yuqiong Guo

## Abstract

The high-quality genome assembly of the JSU-FSP provides valuable information on how jasmine produces the abundant VTs and VPBs that synergistically contribute to the pleasant aroma of jasmine flower.

## Introduction

Jasmine (*Jasminum sambac*) is a tropical or subtropical plant that is notable for the sweet, heady fragrance of its flowers. The high accumulation of volatile terpenes (VTs) and volatile phenylpropanoid/benzenoids (VPBs) in jasmine flowers creates a natural source of essential oil to meet consumer demand all over the world. Additionally, jasmine is also used to produce scented tea, which is reprocessed from finished tea by absorbing aroma volatiles of the fresh flowers and has been favoured since antiquity [1]. Cultivar jasmine varieties mainly belong to three major groups: *J. sambac* var. *unifoliatum* (JSU, single-petal type), *J. sambac* var. *bifoliatum* (JSB, double-petal type), and *J. sambac* var. *trifoliatum* (JST, multi-petal type). The flowers of JSU, with the best aroma quality, are considered as the optimal materials for the production of high-grade jasmine tea [2]. However, the lack of a reference genome limits the studies of basic and applied biology on the jasmine. The genetic bases for the rich production of VTs and VPBs remain largely unclear. While recently the reference genomes of JSB [3] and JST [4] cultivars were *de novo* sequenced and assembled, no JSU genome has yet been reported.

## Results and discussions

Here, we assembled a reference genome for the autotriploid JSU cultivar ‘Fuzhou single-petal’ (JSU-FSP). According to the 21-mer analysis of 45.14 Gb Illumina reads and flow cytometry, we estimated that the haploid genome size of JSU-FSP is ~471.23 Mb and ~498.35 Mb, respectively. We initially assembled the genome by hifiasm v0.16 with 85.89 Gb PacBio HiFi reads, resulting in a 495.60 Mb sequence assembly with a contig N50 of 16.88 Mb. The BUSCO and CEGMA analyses showed that the completeness of the JSU-FSP genome was 97.83% and 98.91%, respectively, which indicated that the excellent continuity and completeness of the assembly. Then, all the contigs were corrected and orientated by Hi-C onto 13 pseudo-chromosomes, generating chromosome-scale sequences of 476.92 Mb (Fig. 1A and 1B).

**Figure 1.**
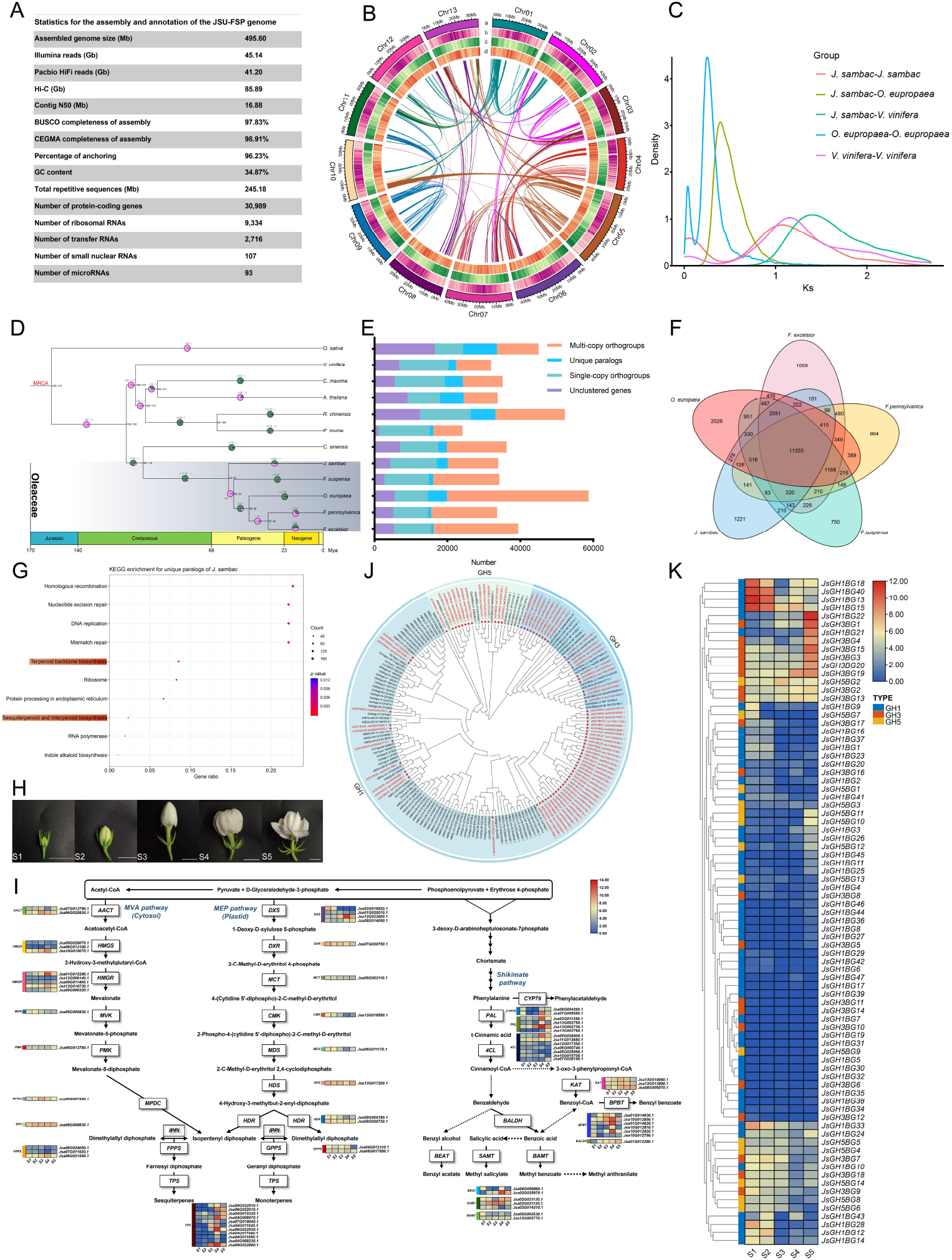
Landscape of the genome of *J. sambac* var. *unifoliatum* cv. Fuzhou single-petal (JSU-FSP). A, statistics for the assembly and annotation of the JSU-FSP genome. B, the circulars map shows chromosome ideograms (a), transposable element density (b), gene density (c), GC content (d), and syntenic blocks (colored lines in the center). C, density distribution of *Ks* within *J. sambac*, *O. europaea*, and *V. vinifera*. D, phylogenetic tree with estimated divergence time and gene family expansion/contraction. E, statistics for the gene family. F, venn diagram of orthogroups within five Oleaceae plants. G, KEGG enrichment analysis for the unique paralogs in JSU-FSP genome. H, images of five flowering stages of JSU-FSP, bar = 1 cm. I, differentially expressed genes involved in terpenes and phenylpropanoid/benzenoids metabolic pathways. J, phylogenetic tree of β-glucosidase (BGLU) proteins from JSU-FSP, *A. thaliana*, and *O. sativa*. K, the expression profile of BGLU genes in different flowering stages of JSU-FSP.

The annotation results showed that there were 264.98 Mb repetitive sequences in JSU-FSP genome, of which long terminal repeat-retrotransposons (LTR-RTs) accounted for 66.33% (175.77 Mb). The Ty3-*gypsy* were the predominant LTR-RTs (58.92 Mb), followed by Ty1-*copia* (36.68 Mb). We identified 30,989 protein-coding genes through integrated *ab initio* modeling, homology-based searches, and transcriptome analysis (Fig. 1A). The BUSCO analysis showed that 96.16% of the core plant genes were present in identified genes, indicating an excellent completeness of genes prediction.

Through comparative genomics analysis between the JSU-FSP genome and the genomes of other 11 representative plant species, we observed that JSU-FSP genome went through an ancient WGD event by estimating *Ks* distributions (Fig. 1C). The phylogenetic tree based on 448 single-copy orthologous genes illustrated that *J. sambac* diverged from the ancestors of other four species of Oleaceae plants about 54~58 million years ago (Fig. 1D). Gene family analysis revealed that 26,638 genes in JSU-FSP were clustered into 17,446 families (Fig. 1E). The five Oleaceae plants shared 11,333 common gene families, while 1,221 gene families were unique in the JSU-FSP genome (Fig. 1F). The unique genes of JSU-FSP were clustered into “Terpenoid backbone biosynthesis” and “Sesquiterpenoid and triterpenoid biosynthesis” of the KEGG enrichment pathways, endowing jasmine with the genetic basis of heady fragrance (Fig. 1G).

As aroma volatiles of jasmine are synthesized and emitted in the opening process [5], flower buds or flowers at five different stages were selected for transcriptome sequencing (Fig. 1H). The expression of key genes was investigated to unveil the mechanism of VTs and VPBs biosynthesis. We identified that among the 73 structural genes involved in VTs and VPBs metabolic pathways, the expression of 43 genes in opening flowers was significantly higher than that in buds. This indicated the differential expression of these related genes in opening flowers was responsible for the aroma formation. In addition to *de novo* synthesis, plant volatiles are present in glycosidically bound forms and can be hydrolyzed to free volatiles by GH1, GH3, and GH5 families of β-glucosidase (BGLU) [6]. Phylogenetic analysis reveled that a total of 81 BGLU genes in JSU-FSP genome were clustered into GH1 (47), GH3 (20), and GH5 (14) subfamilies (Fig. 1J). Among which, 11 BGLU genes were highly expressed in opening flowers, implying that jasmine aroma is also affected by the expression of BGLU genes [7].

## Acknowledgements

This work was supported by the Construction of Plateau Discipline of Fujian Province (102/71201801101) and the Construction Project for Technological Innovation and Service System of Tea Industry Chain of Fujian Agriculture and Forestry University (K1520005A01). We thank Xin Huang from the School of Foreign Languages of Fuzhou University for her linguistic assistance; Ying Yu from Biomarker Technologies for assistance with sequencing; Prof. Yong Li from Henan Agricultural University for providing the genomic data of *Forsythia suspensa*; and Tianlong Fu from Fujian Chunlun Group Co. LTD for helping with material collection.

## Data available statement

Raw sequencing data have been submitted to the NCBI under project number PRJNA795621.

